# Gene regulatory network integration with multi-omics data enhances survival predictions in cancer

**DOI:** 10.1101/2024.12.19.629344

**Authors:** Romana T. Pop, Ping-Han Hsieh, Tatiana Belova, Anthony Mathelier, Marieke L. Kuijjer

## Abstract

The emergence of high-throughput omics technologies has resulted in their wide application to cancer studies, greatly increasing our understanding of the disruptions occurring at different molecular levels. The role of gene regulation as a core driver of biological processes and its role in the development and progression of cancer has been well established. Transcriptional regulation, a crucial aspect of gene regulation, can be represented as a network of interactions between regulators (such as transcription factors) and their target genes. These networks are known as gene regulatory networks (GRNs). Several joint dimensionality reduction (JDR) methods and tools for integrating multi-modal data have been developed in recent years. This study presents a new approach to integrate multi-modal data of different dimensions to consider sample-specific GRNs with multi-omics data in ten cancer datasets from The Cancer Genome Atlas (TCGA). We compare JDR models that include GRNs with those that do not, assessing their association with patient survival. We find that including GRNs can improve associations with patient survival across several cancer types. Focusing on liver cancer, our methodology identifies potential mechanisms of gene regulatory dysregulation associated with cancer progression.

## 1 Introduction

Cancer is a complex and heterogeneous disease caused by disruptions in normal cellular functions. In their seminal papers, Hanahan and Weinberg [1–3] described and characterised the hallmarks of cancer, highlighting the aberrations taking place at multiple molecular levels and the importance of the interplay between them.

The emergence of next-generation sequencing and other high-throughput omics technologies has led to the integration of multiple molecular aspects into cancer studies, increasing our understanding of the disruptions taking place at the different molecular levels. Public databases such as The Cancer Genome Atlas (TCGA) or the International Cancer Genome Consortium (ICGC) are host to libraries of omics data across cancer types. With the availability of these data came a push towards integrative approaches to analysis.

The integration of omics data originating from the same samples has been termed vertical integration [4]. There are three main approaches to vertical integration (reviewed in [5]). The most straightforward approach is known as late integration, which involves analyzing each omics layer separately and identifying correlations and overlaps afterwards. However, this does not account for potential interactions between different molecular layers captured through multi-omics data.

Another approach, early integration, involves concatenating all omics layers into one large matrix, which is then used for analysis. This approach allows capturing interactions between different omics layers; however, the large matrices can be difficult to analyse. Furthermore, this approach ignores differences in distributions across the various omics layers, which can sometimes lead to uninformative patterns being found.

Finally, intermediate integration allows the joint analysis of multi-omics data without concatenating, typically by reconstructing the data in a latent space. This approach addresses the main drawbacks of the previous two. Intermediate approaches generally involve dimensionality reduction and have thus been termed joint dimensionality reduction (JDR) approaches [6]. Many intermediate integration tools have been published in recent years [6, 7], including MOFA+ [8], JIVE [9], MCIA [10], and RGCCA [11]. One challenge for JDR methods is the difficulty reconciling data types whose dimensionalities differ by orders of magnitude. Since dimensionality reduction is being performed jointly on multiple data types, differences in their dimensionalities can cause the latent space to be mostly influenced by the data types with higher dimensionalities, leading to potential bias.

The role of gene regulation as a core driver of biological processes and its role in the development and progression of cancer has been well established [12, 13]. Therefore, understanding gene regulation and how it is disrupted in diseases is one of the key challenges in modern medical molecular biology. Gene regulation is a complex process encompassing multiple layers of controls and molecular interactions. An essential aspect of gene regulation is transcriptional regulation, which involves interactions between regulators and their target genes. These interactions can be represented as gene regulatory networks (GRNs) [14]. One such interaction is between transcription factors (TFs), which bind promoters and enhancers in a sequence-specific manner to facilitate transcriptional regulation, and their target genes. Numerous tools have been developed for GRN inference both from bulk [15, 16] and single-cell [15, 17, 18] data.

Among the tools developed for bulk data is PANDA (Passing Attributes between Networks for Data Assimilation) [19]. PANDA’s core method is based on the assumption that TFs that cooperate are likely to share target genes, and target genes that are co-expressed are likely to be regulated by similar sets of TFs. The method thus uses a message-passing approach to find agreement between three networks: a TF protein-protein interaction prior, a TF sequence-based prior, and gene co-expression. The output of PANDA is a genome-wide, population-level “aggregate” GRN. However, in the context of cancers, it is important to model GRNs for individual patients. LIONESS (Linear Interpolation to Obtain Network Estimates for Single Samples) [20] can be used to calculate sample specific GRNs based on an aggregate network.

Because of the important role transcriptional regulation plays in cancer, we aimed to assess whether single-sample GRNs modelled with PANDA and LIONESS are informative of cancer patient survival when integrated with other omics data (Figure 1). We benchmarked a data-driven filtering approach to address the issue of dimensionality differences in omics when considering four JDR approaches. We then performed JDR on multi-omics data for ten cancer types from TCGA and compared the models we obtained without the GRNs to those incorporating GRNs, investigating various network metrics. We found that overall, GRNs boost associations with survival. Focusing on liver cancer, we found that considering GRNs led to a drastic increase in association with survival. These differences were driven by increased regulation of metabolic pathways and TFs involved in immune signalling. These findings were replicated in a curated collection of independent liver cancer data. This emphasizes the role of dysregulated metabolic signalling in liver cancer prognosis. Altogether, our results highlight the importance of considering GRN features in multi-omics analyses to advance our understanding of cancer mechanisms at the individual patient level, which could be leveraged for the development of personalized therapies in the future.

**Figure 1.**
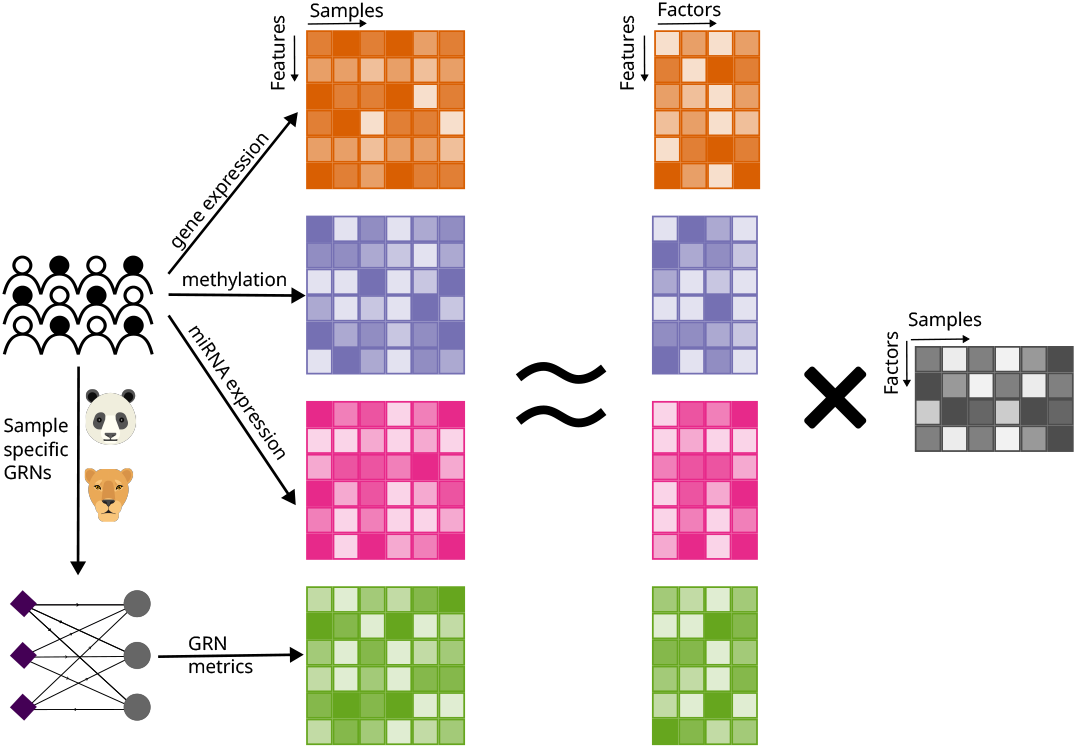
Overview of approach; 10 multi-omics cancer datasets from TCGA were integrated with PANDA/LIONESS sample-specific gene regulatory network metrics using joint dimensionality reduction.

## 2 Methods

### 2.1 Data acquisition and processing

The TCGA data were downloaded from and processed as described in the MOMIX benchmarking pipeline [6]. The data consisted of RNA-sequencing (RNA-seq), microRNA (miRNA)-seq, and methylation array data. Several filtering steps were performed on both the samples and the omics features to ensure all methods could run. The features with *sd* = 0 were filtered out. The GRNs were constructed on the complete set of samples downloaded from MOMIX. For JDR, only samples with data available for all omics types were considered as it is a requirement for some of the JDR methods tested. Finally, for the survival analysis, we only considered samples without missing values in the clinical data. A summary of the different datasets and the number of samples used in our analyses can be found in Table S1.

Additionally, bulk RNA-seq data for 376 liver tumour samples was downloaded from GEPliver [21] along with clinical data. We filtered features and samples as described above.

### 2.2 Gene regulatory networks

Sample-specific GRNs were generated with PANDA [19] and LIONESS [20] using the MATLAB implementation of both tools, as in [22]. Briefly, PANDA uses a message-passing approach to update a prior network of potential interactions with gene co-expression and protein-protein interaction data. The prior network is based on a TF motif scan in gene promoters. PANDA outputs a complete, bi-partite aggregate GRN with weights representing the likelihoods of interactions (see section 2.3). LIONESS uses linear interpolation to infer sample-specific GRNs from an aggregate GRN (e.g. constructed by PANDA). By assuming that each sample linearly contributes to the aggregate network, LIONESS iteratively excludes one sample and reconstructs the aggregate network without it. The sample-specific GRN corresponding to the excluded sample is built from the difference between the aggregate GRN and the newly reconstructed one, corrected for sample size. Sample-specific GRNs were constructed independently for ten cancer types in TCGA and for the GEPLiver dataset.

### 2.3 Bi-partite networks and network metrics

The resulting GRNs are bi-partite with nodes corresponding to TFs and target genes and directed edges only existing between these two types of nodes, flowing from TFs to genes (Figure S1A). These networks are complete, with all possible TF-gene edges represented in the graph. Edge weights in this graph represent the likelihood that a particular TF-gene interaction occurs. Edge weights can be negative, however the sign does not indicate direction of regulation, lower weights simply representing lower likelihood interactions.

We used the network degree to summarise the GRNs for use with JDR. Since our networks are bi-partite, we calculated the indegrees and the outdegrees. The *indegree* (Figure S1B) of a gene is the sum of all the edges (or edge weights) that connect it to its regulators; it measures how much regulation each gene receives from all its regulators (i.e., one would expect genes that are more highly regulated or that perhaps have more redundancy in their regulation, to have higher indegrees). The *outdegree* (Figure S1C) of a TF is the sum of all edges (or edge weights) that connect it to its targets in a GRN; it measures how much a TF regulates its targets (i.e., one would expect more active TFs to have higher outdegrees). It is important to note that the degree cannot inform of the directionality of the regulation (i.e., activation versus repression).

### 2.4 Benchmarking PCA versus no PCA

We used a modified version of the benchmarking described in [6]. We followed the same general protocol but used the most recent versions of all tools in R version 4.2.1. Furthermore, since the result of the original benchmarking indicated that the performance of the tools was dependent on the data type, we selected four of the tools that implement linear decomposition of the omics matrices and that performed well on the TCGA dataset in the original benchmarking study (JIVE, MOFA, MCIA, and RGCCA). A description of each algorithm can be found in [6]. While the original benchmark used MOFA, we used MOFA+, described as an improved version in [8].

PCA was performed in R with the *pcamethods* package using the single value decomposition (SVD) approach. For each data type, a cumulative *R*^2^ score of 0.85 was used to determine the number of PCs for the factorisation. We also enforced a minimum of 20 PCs in cases where fewer were needed to reach the variance threshold.

Although each method can optimise the number of factors for itself, we imposed five factors for all methods to perform the comparisons. We acknowledge that, in some cases, some of the factors identified explained virtually no variability. Some JDR tools, such as MOFA+ discard such factors by default.

### 2.5 MOFA+ models

MOFA+ was run with default parameters, except for (i) the number of factors set to five, (ii) the convergence mode set to *slow*, and (iii) the seed set to 13. We randomly selected five seeds and performed the survival analysis (see section 2.7) for each to ensure that the choice of seed has no impact on the results since MOFA+ is a stochastic algorithm. We did not observe any significant differences in results across the different seeds (Table S2). Thus, we report the results obtained with our original seed (13).

Separate models were run for the omics data without the networks, with the network indegrees, with the network outdegrees, and with both indegrees and outdegrees.

### 2.6 Mapping back to original features

One of the advantages of JDR tools such as MOFA+ is the ability to determine what features (e.g., genes) are important to each factor (i.e., feature weights). However, since we performed PCA on the data before JDR and used principal components as input into MOFA+, we lost the direct mapping of the factors to the features. Instead, we have a mapping of factors to principal components (MOFA+ weights) and a mapping of principal components to the original features (PCA loadings). We, therefore, multiplied the matrix of PCA loadings to the matrix of MOFA+ factors to obtain an approximate mapping of the MOFA+ factors to the original omics features. For a mathematical explanation, see Supplementary methods.

### 2.7 Association with survival and clinical features

The association of latent factors with patient survival was performed using the *survival* package (version 3.3.1) in R. For each factor, we fit a univariate Cox proportional hazards model. For the association with other clinical features, we performed Wilcoxon rank-sum (Mann-Whitney) tests for binary features and Kruskal-Wallis tests for other features. We performed Benjamini-Hochberg false discovery rate correction on the p-values.

### 2.8 Gene set enrichment analysis

Gene set enrichment analysis (GSEA) was performed using the *fgsea* function from the *fgsea* R package using the MSigDb [23, 24] Hallmark [25], and KEGG-legacy [26] gene sets obtained with *msigdbr* [27]. We used the feature weights (see section 2.6) as rankings.

### 2.9 Outdegree analysis

We assessed the overlap in the top outdegree features for the survival-associated factors (SAFs) in the TCGA liver and GEPliver datasets. First, we selected the top 20 TFs per SAF with the highest absolute weights. We then performed a Fisher’s exact test to determine if the overlap between the two sets was larger than expected by chance.

## 3 Results

### 3.1 PCA as a data-driven approach to feature-selection for JDR

One of the challenges of multi-omics integration is the different characteristics of the various omics data types. For example, RNA-seq data and methylation array data represent different kinds of signals with different distributions, and they have numbers of features that differ by orders of magnitude. Finding a statistically robust way of bringing such different data together and preserving the underlying biological signals remains challenging.

Due to the large variation in the number of features captured by different omics, most JDR methods recommend that the user filter the data so that the omics layers have comparable dimensions. This is recommended to avoid the results being driven by the difference in dimensionality between omics layers, rather than by biological signal. Filtering is typically done by selecting the *n* most variable features, *n* often being 5000, which may not always capture the most relevant variability. Furthermore, not all omics are made equal; the top 5000 features in gene expression and methylation data for example, could represent a very different proportion of the overall variability in their respective datasets. Another commonly used but more relaxed filtering approach is to remove non-variable features. However, this approach may not solve the issue of dimensionality differences.

One data-driven approach to filtering is to perform principal component analysis (PCA) on the data before JDR, using a variance threshold to filter the data. We tested several thresholds and found that 85% of the variability was a suitable compromise between capturing as much variability as possible and keeping the omics dimensions with one order of magnitude of each other. However, the dimensionality gap reduced at even higher thresholds (Figure S2). The numbers of principal components (PCs) capturing 85% of the variability exhibited less variation than the original number of features (Figure 2A, Figure S3). Most JDR methods are robust to differences in omic dimensions within an order of magnitude. We found that performing PCA before JDR and using a cumulative *R*^2^ threshold to select the number of PCs consistently yielded dimensions within an order of magnitude of each other across cancer datasets (Figure S3).

**Figure 2.**
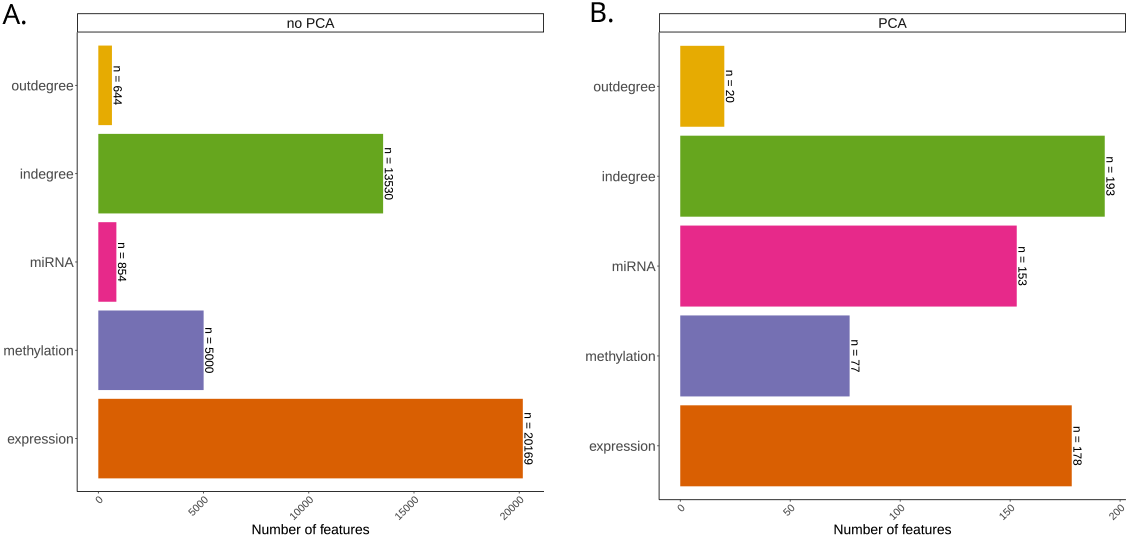
Comparison of data properties before and after PCA in liver cancer. **A**. Number of features before PCA. **B**. Number of selected PCs. Results for all tested cancer types are shown in Figure S2.

However, using PCA has its own limitations. First, as dimensionality reduction is performed, some information is lost, albeit in a more data-driven fashion. Second, it adds a layer of complexity to the interpretation of the results (see section 2.6).

To determine whether performing PCA on the data before JDR is viable, we assessed the association of the resulting factors with patient survival in models obtained with the original features and with data where PCA was performed before applying JDR. We followed the benchmarking pipeline described in [6] (see section 2.4). We found that the performance varied between the datasets and the JDR methods (Figure S4). For MOFA, MCIA and JIVE, the performance was, in general, better when PCA was used. When considering the top factor for each cancer to assess the sensitivity of the models, we found that MOFA and MCIA performed better in 7/10 datasets and JIVE in 5/10. The exception was RGCCA, which performed considerably worse in all datasets when using PCA.

We opted to use MOFA+ for the remainder of the study as it offered the best balance. JIVE was slower to run than any other methods, while MOFA+ was the only tool to allow some samples to be missing from some omics types, a valuable feature for future applications. However, for this analysis, we considered samples with observations for all omics types to allow for comparison between methods.

### 3.2 GRNs better identify factors associated with survival

Next, we assessed whether GRNs are informative of patient survival in the context of JDR. We ran MOFA+ separately on all omics data types with no GRN information and compared the results with MOFA+ models that included network metrics (indegree, outdegree, and both).

When we assessed the association between the resulting factors and patient survival, we found that adding the indegrees had little impact in most cancer datasets. However, in the case of kidney cancer, the consideration of indegrees decreased the significance of the association (Figure 3A). Considering outdegrees revealed a factor associated with survival in the AML dataset, while no such factor was found with the omics-only model or the model considering indegrees (Figure 3B). However, it did not reveal relevant factors for the remainder of the cancer types.

**Figure 3.**
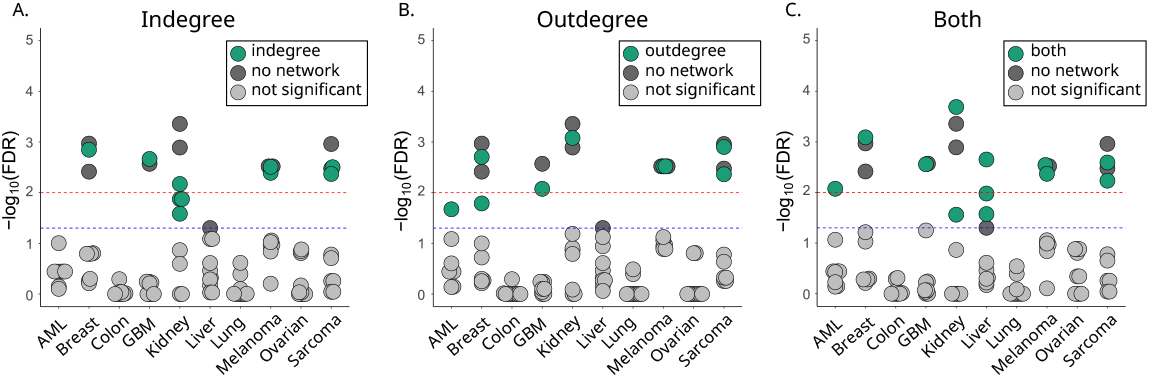
Comparison of factor association with patient survival between MOFA+ models without networks and MOFA+ models with network indegrees (A), outdegrees (B) or both (C). Each dot represents the association of one factor with patient survival.

Finally, adding indegree and outdegree features to the model revealed factors significantly associated with survival in the AML, kidney, and liver datasets. The association of these factors with survival was more significant than factors obtained with the omics-only models. The analysis of the liver dataset yielded three survival-associated factors that we did not find with any of the previous models, likely highlighting the interplay between different omics layers (Figure 3C).

### 3.3 GRN features integration with multi-omics data reveals survival-associated factors in liver cancer

We explored the survival-associated factors (SAFs) identified when including GRN features for the liver cancer samples (Figure 4A). We first evaluated which feature types were explained by the SAFs (Figure 4B). For all three SAFs, the MOFA+ model explained the most variance in the expression data, followed by miRNA, indegrees and outdegrees. Notably, despite the comparatively small percentage of variance explained in the networks, their inclusion was crucial to revealing these SAFs. One possibility is that the GRNs are simply boosting a signal captured by some of the other omics layers.

**Figure 4.**
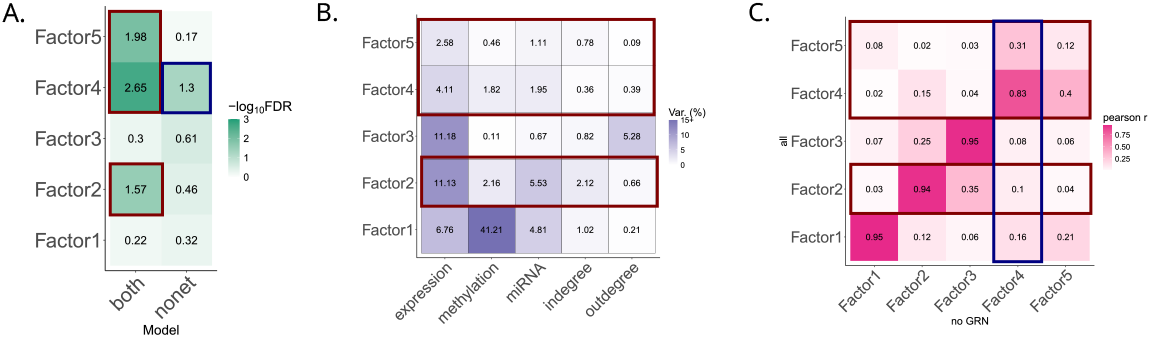
Exploring MOFA+ factors in liver cancer. Highlighted in red are the three SAF in the network model and in blue the SAF found in the model without networks. **A**. Heatmap of −*log*_10_(*FDR*) of MOFA+ factor association with patient survival in models with (“both”) and without (“nonet”) networks. **B**. Heatmap showing the percentage of variance explained by the MOFA+ factors in each omics type and network metric. **C**. Heatmap showing the pairwise absolute Pearson correlation between MOFA+ factors from the model including the GRN metrics and the model without any GRN metrics.

To investigate this possibility, we performed pairwise correlation between the factors from the omics-only model and the model including the GRN features (Figure 4C). We found that Factors 2 and 4 from the models including the GRNs correlated well (pearson r ≥ 0.8) with Factors 2 and 4, respectively, from the omics-only models. In the case of Factor 2, it confirms that adding GRN features boosts a signal found in the other data modalities (Figure 4B-C). Indeed, Factor 2 explains variability in the mRNA and miRNA modalities (Figure 4B).

Factor 4 was the only factor associated with survival in the omics-only model, showing that we do not “lose” any SAFs with the addition of the GRN features despite maintaining the five factor constraint. Finally, Factor 5 did not correlate well with any of the factors in the omics-only model, suggesting that the GRNs may capture some heterogeneity within the population that is not present, or is very weak in the other omics modalities.

### 3.4 Independent validation of multi-omic association with survival in liver cancer

We performed the same analysis in an orthogonal liver cancer dataset from GEP liver [21], which comprised ∼370 samples profiled with RNA-seq and had clinical annotation available (Table S2). As with the TCGA liver dataset, we identified three factors associated with patient survival (Table S3). We evaluated the association of the MOFA+ factors with other clinical features (Figure S5) in both datasets. Age was not significantly associated with any of the factors in either dataset. Liver fibrosis was associated with the three SAFs in the TCGA dataset and two of the three SAFs in the GEP dataset. Histological subtype was associated with all SAFs in both datasets, however the TCGA dataset is comprised predominantly of hepatocellular carcinoma (HCC), with a very small number of samples representing other subtypes, while the GEP dataset is more heterogeneous.

### 3.5 Increased regulation of metabolic pathways is associated with liver cancer survival

To determine the contribution of GRN features in multi-omic integrative approaches, we further looked at what drives these SAFs in the indegree and outdegree spaces. We performed GSEA on the weights of the indegrees for each of the SAFs in both the TCGA and GEP datasets using the MSigDb Hallmarks [25] gene sets (Figure 5A-B, Figure S6A-B).

**Figure 5.**
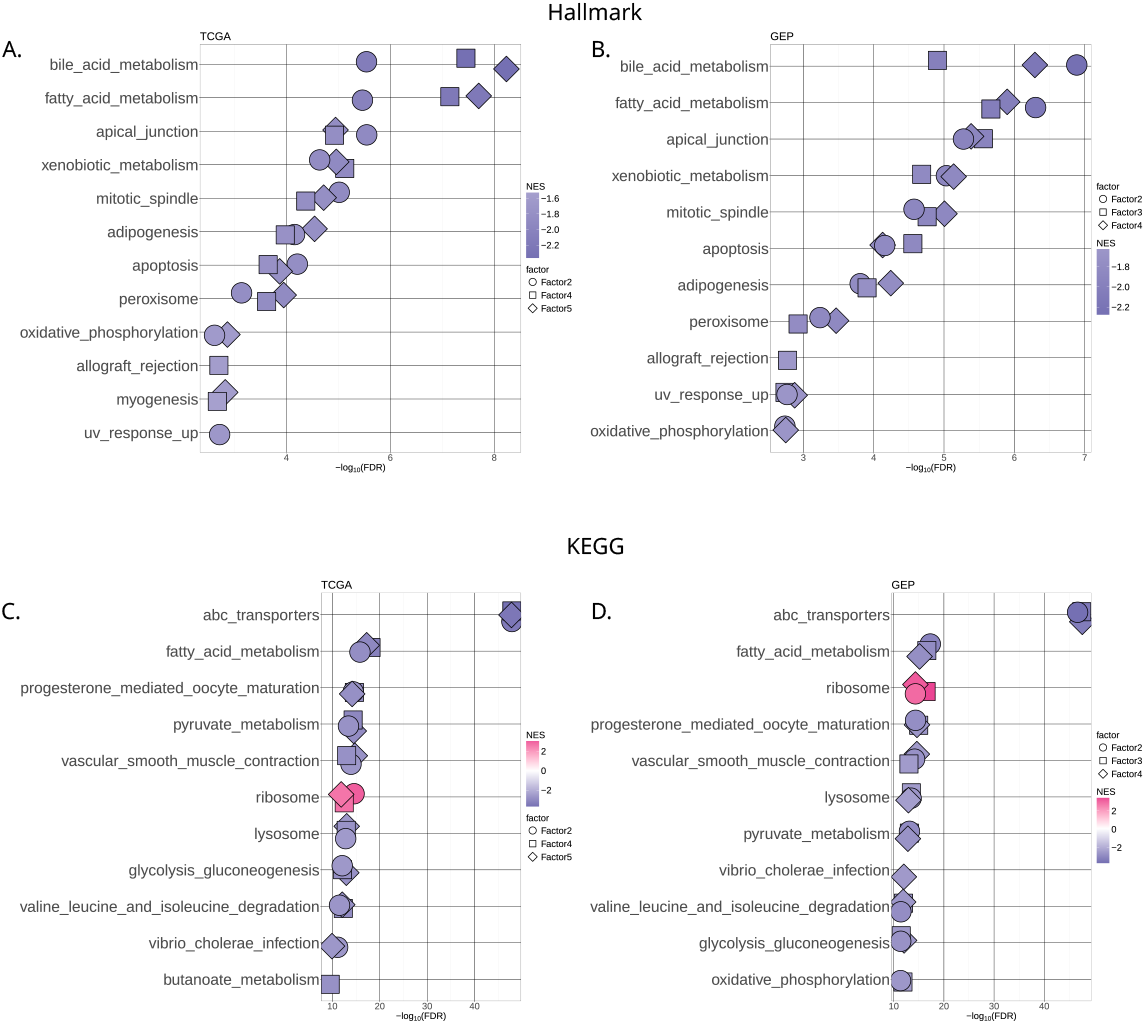
GSEA on the MOFA+ indegree weights of SAFs in the TCGA datasets using the hallmark (**A-B**) and KEGG (**C-D**) gene sets.

The enriched pathways are consistent across the two datasets. Prominent among them are several metabolic pathways. Studies have found reprogramming of metabolic pathways to be strongly connected with prognosis in primary liver cancer [28, 29]. To get a more granular view of the metabolic pathways driving the SAFs in the indegree space, we performed enrichment analysis of KEGG [26] gene sets (Figure 5C-D, Figure S6C-D). Fatty acid metabolism was one of the top pathways in both the MSigDb Hallmarks and KEGG analyses. Along with glucose and amino-acid metabolism, lipid metabolism was found to be one of the main metabolic alterations in HCC. [29, 30]. Both biosynthesis and degradation of fatty acids have been shown to play a role in liver tumour progression. Degradation of fatty is often down-regulated, while de-novo synthesis of fatty acids is often up-regulated. Indeed, the fatty acid degradation pathway classifies liver tumours into three molecular subtypes, which, in addition to being indicative of prognosis, responded differently to various treatments [31].

In addition to fatty acid metabolism, ABC transporters pathways were strongly enriched in both datasets. A growing body of literature (reviewed in [32]) links ABC transporters and cancers. A few studies have linked ABC family members to liver cancer [33, 34], showing their involvement in several metabolic and inflammatory processes.

Most of the top pathways show a negative enrichment in both datasets. MOFA+ factors ordinate the samples along an axis centered at 0, with the signs indicating the directionality of the phenotype (i.e., samples with opposite signed factor values have opposite phenotypes), and the magnitude representing the strength of the phenotype. The feature weights score the relationship between each feature and the phenotype captured by each factor. A positive weight means a feature has higher levels in samples with positive factor values. A negative sign means a feature has high levels in samples with negative factor values [8]. Thus, negative enrichment suggests that samples with low (negative) factor values have high indegree values for genes in the enriched pathways. When splitting the cohorts based on the median of the SAFs and plotting Kaplan-Meier curves of the resulting groups (Figure S7), we see that the samples with low factor values correspond to better survival. Therefore, stronger regulation of these pathways associates with improved outcomes.

### 3.6 Immune regulatory and developmental TFs associate with liver cancer survival

To assess the contribution of the outdegrees, we opted to look at the top TFs driving the SAFs in each of the two datasets. We selected the top TFs based on MOFA+ weights for each SAF. Twenty-three TFs were shared between the two datasets. Due to differences in the composition of the two datasets, using a threshold on the weights to select the top TFs yielded different number of TFs, leading to an overlap that was not significant (fisher’s exact test odds ratio 0.66, p-value 0.2).

We observed that several shared TFs were ranked at the top (i.e., had high absolute weight values in at least one SAF in each dataset) in both datasets (Figure S8). We postulated that the greater heterogeneity of the GEP dataset could be a confounding factor but that the same heterogeneity as in the TCGA dataset may be present in the HCC subpopulation of the GEP dataset (which is also the predominant subtype in that data). We decided to narrow our focus to the top of the TF rankings and enforce a similar number of TFs for the two datasets.

We selected the top twenty TFs with the highest weights for each SAF in each dataset and looked at the TF overlap between the two datasets. We found a significant (Fisher’s exact test odds ratio 2.68, p-value 0.02) overlap of nine TFs (Figure 6). We find several TFs involved in immune regulation (RFX1, RFX5, JUND). RFX1 has been identified as a potential therapeutic target for cancers due to its involvement in many processes that are thought to interfere with cancer stem cells [35]. In addition, several developmental TFs are present (HOXB1, ARID2, TBX15, MESP1). RBPJ is involved in Notch signalling and is an activator of the pathway’s downstream targets, however it can also act as a repressor when not associated with Notch.

**Figure 6.**
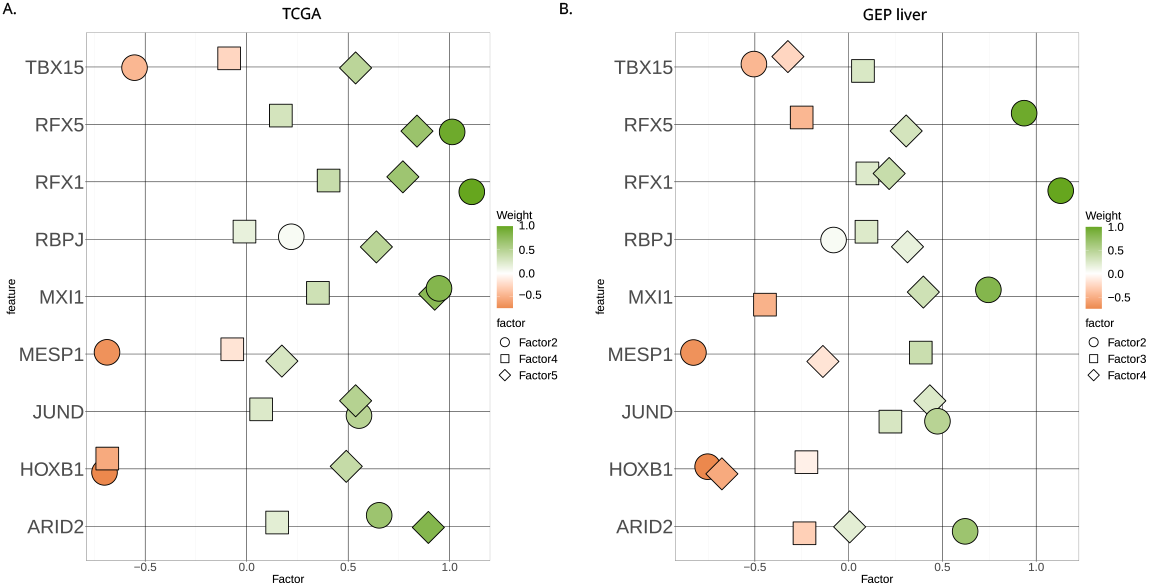
Weights of common TFs in TCGA liver (**A**) and GEP liver (**B**) datasets.

Together, these results highlight the dysregulated signaling pathways in liver cancer progression and underscore the value of multi-omics integration with gene regulatory networks in individual cancer patients.

## 4 Discussion

In this study, we explored PCA as a data-driven approach to filtering multimodal data for joint dimensionality reduction methods, which can allow the application of JDR on data types with different dimensions. We leveraged this approach to investigate single-sample GRNs in the context of multi-modal data integration. We found that GRNs can improve the sensitivity of JDR models to identifying survival-associated factors.

JDR methods can integrate multi-modal data by reducing them to a shared latent space (i.e., latent factors). However, one limitation of these approaches is that they are not robust to large differences in dimensionalities of the different data modalities. While they can generally accommodate some variation in the number of features, it is generally advised to keep them within an order of magnitude of each other, with some tools such as MOFA+ [8] explicitly making this recommendation.

We explored PCA as an alternative, more data-driven approach to solving this issue. Instead of imposing a threshold on the number of features, we performed PCA on each omics data type and then used a variance threshold to select the number of PCs. We found that this brought the different data types closer regarding the number of features while representing a similar proportion of the overall variability of the respective datasets. Furthermore, as PCs represent a combination of all the features in a dataset, this approach allowed all features to contribute to the downstream JDR, unlike a hard cutoff on the number of features.

One obvious limitation of this approach is the loss of the mapping between the latent factors obtained with JDR and the omics features, limiting the downstream analysis options available. However, we show that it is possible to reconstruct an approximation of this mapping when using JDR tools that employ linear decomposition (see Supplementary Methods). We do not recommend this approach when using non-linear JDR tools.

We tested our filtering approach on four linear JDR tools: MOFA+, MCIA, RGCCA and JIVE. We specifically focused our benchmarking on identifying factors associated with patient survival, as that was our intended application. In the case of MOFA+, JIVE and MCIA, the performance was either unchanged or improved by the PCA filtering in most of the tested datasets. RGCCA was the only one that performed notably worse across all datasets when using the PCA filtered data.

RGCCA works by first modelling omic specific factors and then maximising the correlation of factors from different omics, to identify joint factors. MCIA has a similar approach, but maximises co-inertaia instead of correlation. When benchmarked, correlation-based approaches like RGCCA performed worse than co-inertia based approaches [6]. It is possible that performing PCA interferes with the correlative relationships between omic-specific factors, thus leading to worse performance.

GRNs have become widely used, versatile tools for representing and studying the complex layers of interactions governing gene regulation [15], and add an additional layer of information to other omics data types, such as gene expression. GRNs are often built on gene expression data, however, it has been previously shown that PANDA-LIONESS GRNs, and in particular the indegrees, generally do not correlate with gene expression [36], suggesting GRNs capture different heterogeneity. However, the question remains whether they would be useful in multi-modal data integration.

We generated single-sample gene regulatory networks for ten cancer datasets from TCGA, calculated the indegrees and outdegrees and used them for multi-omics integration with MOFA+. Comparing the power of MOFA+ models with and without patient-specific GRNs in terms of factor association with patient survival, we found that, in general, the cancer type was the main determinant of whether any SAFs were identified. This is in line with previous findings [6]. Without studying additional datasets, it is difficult to say whether this results from biological heterogeneity between different cancer types or technical heterogeneity between the datasets. Likely, it is a combination of the two.

However, we did observe an increase in the significance of the association with survival when adding indegrees and outdegrees to the MOFA+ models in a few cancer datasets. In AML and liver cancer, we identified additional SAFs not captured by the models without networks. The addition of indegrees alone made little difference. While the addition of the outdegrees alone showed some improvement compared to the model without networks, the addition of both yielded the most striking results.

Finally, we looked closer at the SAFs identified in liver cancer and reproduced the analysis in an orthogonal dataset. Gene set enrichment in the indegree space identified various metabolic pathways enriched, including fatty acid metabolism. Lipid metabolism disruptions have been well documented in liver tumours (reviewed in [29, 30]). A recent study [31] proposed different treatment strategies based on differences in fatty acid metabolism, highlighting its potential as a biomarker for personalised treatment approaches.

The link between lipid metabolism and the immune system in liver tumours has previously been established [37, 38]. Metabolic alterations in the tumour microenvironment can lead to changes in the immune response in the liver. Furthermore, metabolic reprogramming can also occur within local immune cells in the tumour, altering their function and facilitating tumour progression.

We identified several TFs involved in immune regulation as drivers of SAFs, such as JUND, a JUN family TF that forms part of the AP-1 complex. AP-1 is involved in several processes underlying tumour development and progression, serving as both an oncogene and tumour suppressor depending on context and the specific JUN/FOS proteins forming the complex [39]. JUND, as part of the AP-1 complex, has been linked with hepatic lipid metabolism and non-alcoholic fatty liver disease (NAFLD) [40], one of the increasingly common precursors to liver tumours. The role of JUND specifically in liver tumours is still poorly understood, however, our results identify it as a potentially interesting candidate for further research, as it could be a regulatory driver of lipid metabolism.

### 4.1 Conclusions and future perspectives

This study investigated the utility of GRNs in multi-modal data integrative approaches and presented a framework for incorporating them into such analyses. We found that GRNs can be successfully used in such approaches and have the potential to yield insights into the mechanisms of disease. As GRNs model complex interactions and often are themselves an integration of several sources of information (e.g. priors), they can highlight interactions that other omics data types may not easily capture.

## Reproducibility, data and code availability

All data used for this analysis can be found on Zenodo (https://zenodo.org/records/14524447). The code for reproducing the analysis and figures presented here is available on GitHub at [JDRnet](https://github.com/rtpop/JDRnet). A container is available on Zenodo with the software versions used for this analysis.

## Acknowledgments

The authors would like to acknowledge and thank Torfinn Nome for help with creating a container for the reproducibility of the analysis and Ine Bonthuis and Patricio López Sánchez for help with testing code.

