## Supplementary material for "Gene regulatory network integration with multi-omics data enhances survival predictions in cancer"

#### Supplementary methods

##### Mapping factor weights to omics features when using PCA prior to JDR

In order to perform downstream analysis with the MOFA factors, we needed to map the MOFA factors back to the original features (such as genes). We approximated this mapping as:

$$W_{\text{MOFA}} \approx P_{\text{filtered}} W_{\text{PC-fil}} \tag{1}$$

Here, $W_{\text{MOFA}}$ is the matrix of MOFA weights for one omic dataset, $P_{\text{filtered}}$ is the matrix of PCA loadings filtered based on a cumulative $R^2$ threshold (see Section 2.4 in the main manuscript), and $W_{\text{PC-fil}}$ is the matrix of MOFA weights for the filtered principal components of the omic dataset. Below is the mathematical justification.

Given an omic matrix $X$, PCA finds a matrix of PCA scores $S_{\text{PCA}}$ as given by (2) [1]. $P$ is the matrix of PCA loadings.

$$S_{\text{PCA}} = XP \tag{2}$$

From (2), we can express $X$ as (see relationship between PCA and single value decomposition [1, 2]):

$$X = S_{\text{PCA}} P^T \tag{3}$$

Let's now consider the matrix factorization performed by MOFA. Given $Q$ omic matrices $Y_i$, for $i = 1, \ldots, Q$, MOFA implements a linear factorization of the omic matrices as given by (4) [3, 4].

$$Y_i = W_{\text{MOFA},i} F_{\text{MOFA}} + E_{\text{MOFA},i}, \text{ for } i = 1, \ldots, Q \tag{4}$$

Where $F_{\text{MOFA}}$ is a factor matrix common to all omics, and $E_{\text{MOFA},i}$ is the error or residual noise. For simplicity, we will focus here on only one omic matrix $Y$.

$$Y = W_{\text{MOFA}} F_{\text{MOFA}} + E_{\text{MOFA}} \tag{5}$$

One important note is that PCA and MOFA expect input in different formats. PCA expects input to be formatted as samples $\times$ features, whereas MOFA expects input with features $\times$ samples format. Thus:

$$Y = X^T \tag{6}$$

From (3) and (6):

$$Y = PS_{\text{PCA}}^T \tag{7}$$

In our analysis, we input the transposed PCA score matrix $S_{\text{PCA},i}^T$ into MOFA instead of the original omic matrix $Y$.

$$S_{\text{PCA}}^T = W_{\text{PC}}F_{\text{PC}} + E_{\text{PC}} \tag{8}$$

From (7) and (8):

$$Y = P(W_{\text{PC}}F_{\text{PC}} + E_{\text{PC}}) \tag{9}$$

Which we can expand as:

$$Y = PW_{\text{PC}}F_{\text{PC}} + PE_{\text{PC}} \tag{10}$$

Let $PE_{\text{PC}} = E_{\text{combined}}$, we now have:

$$Y = PW_{\text{PC}}F_{\text{PC}} + E_{\text{combined}} \tag{11}$$

We have arrived at a reformulation of the MOFA decomposition. From (5) and (10), we can see that $W_{\text{MOFA}} = PW_{\text{PC}}$. However, this equation holds true when using a complete set of principal components and PCA loadings. Since we filtered the principal components based on a cumulative $R^2$ threshold prior to performing MOFA, we introduce an additional small error, and therefore, this becomes an approximation. Thus $P$ becomes $P_{\text{filtered}}$, $W_{\text{PC}}$ becomes $W_{\text{PC-fil}}$, and we arrive at our starting equation:

$$W_{\text{MOFA}} \approx P_{\text{filtered}}W_{\text{PC-fil}} \tag{1}$$

#### MARMOT: A tool for JDR model comparison and analysis

There are many tools for joint dimensionality reduction of multi-omics data, many of which have been benchmarked [5], however the suitability of various tools seems to be largely dependent on the data and the downstream analysis being performed.

In our analysis, we found the need for comparing JDR models under different conditions. Here we present MARMOT (**M**odel **A**nalysis and compa**R**ison for **M**ulti-**O**mics **T**ools), an R tool for comparing JDR models in different conditions or with different inputs and performing various downstream analyses. MARMOT builds upon the MOMIX benchmarking pipeline published by Cantini et al. [5], with four JDR tools currently implemented.

MARMOT can be accessed and installed from:
`https://github.com/rtpop/MARMOT`

#### Supplementary figures

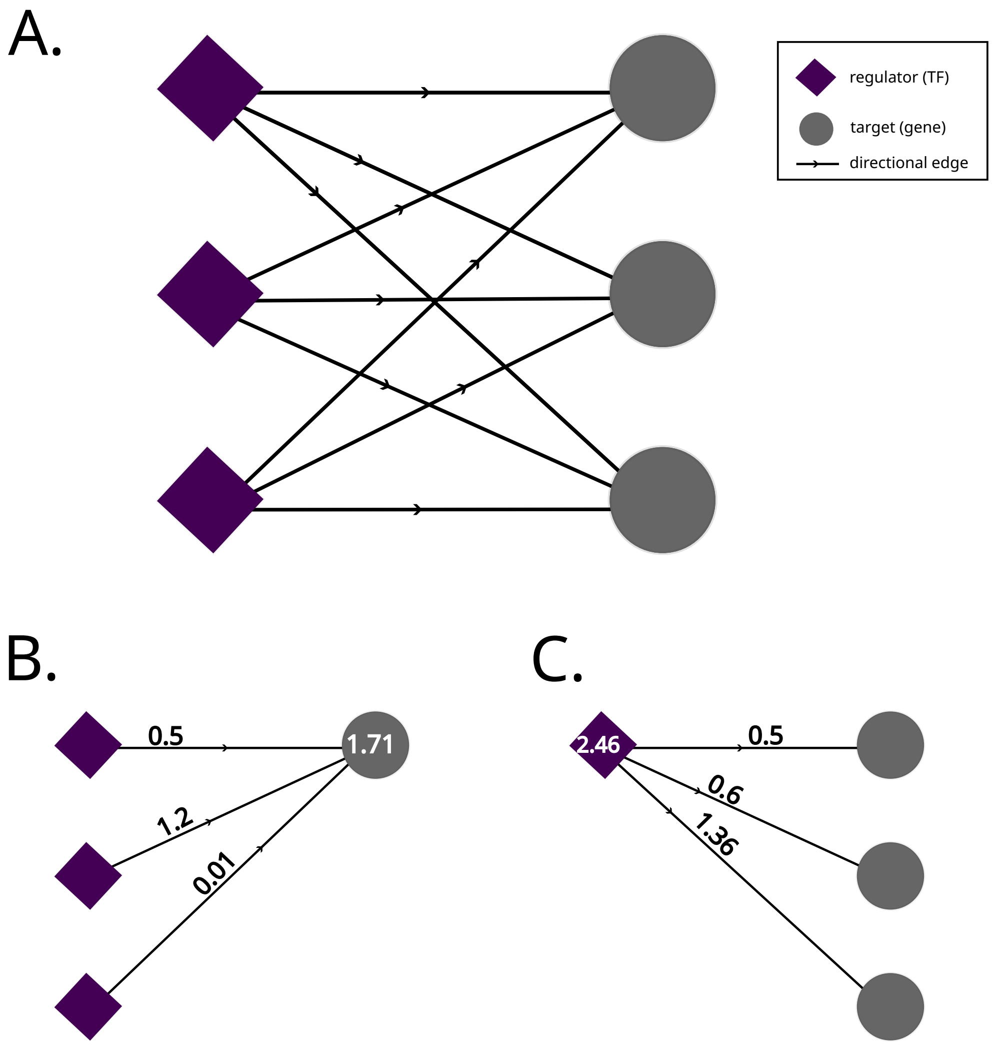


**Figure S1.** **A.** Illustration of a bi-partite, complete and directional network. **B.** Illustration of the indegree calculation. **C.** Illustration of the outdegree calculation.

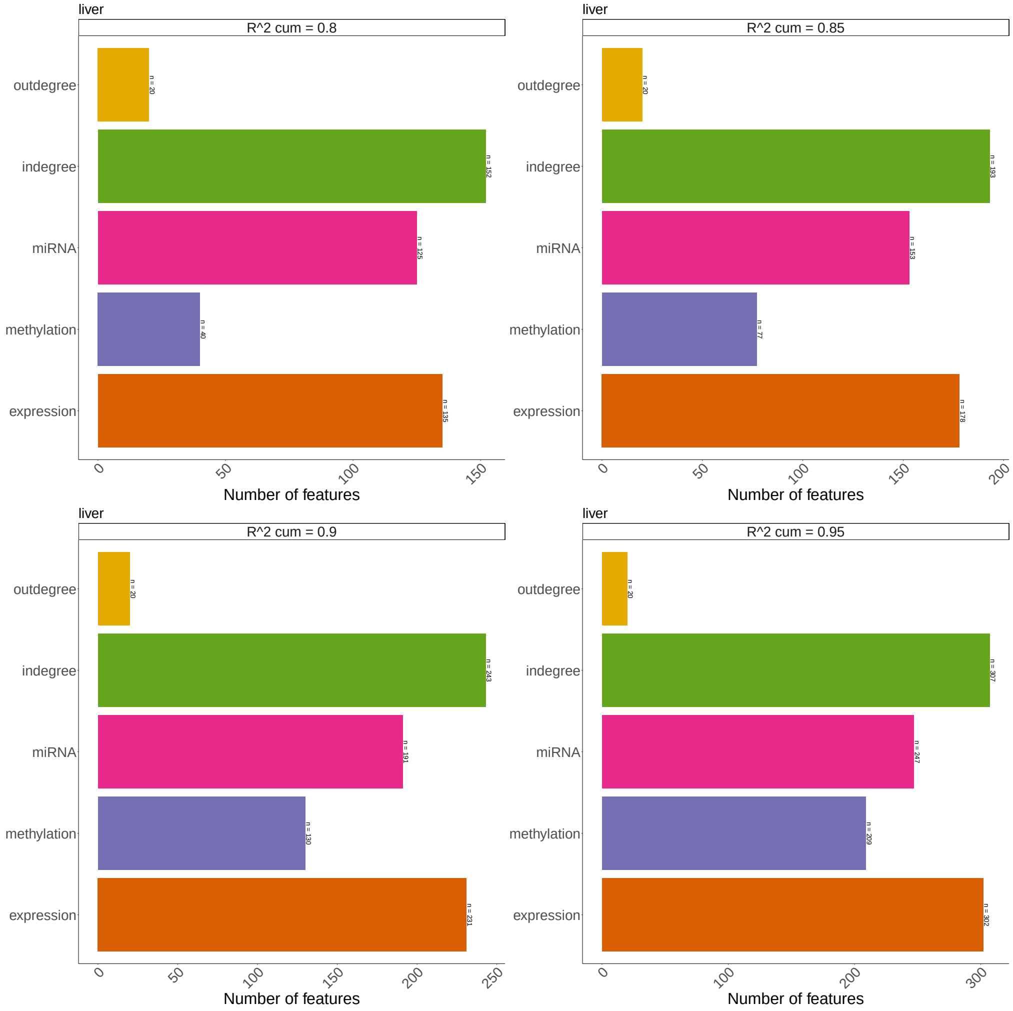

**Figure S2.** PCA data dimensions when using various cummulative $R^2$ thresholds in liver cancer.

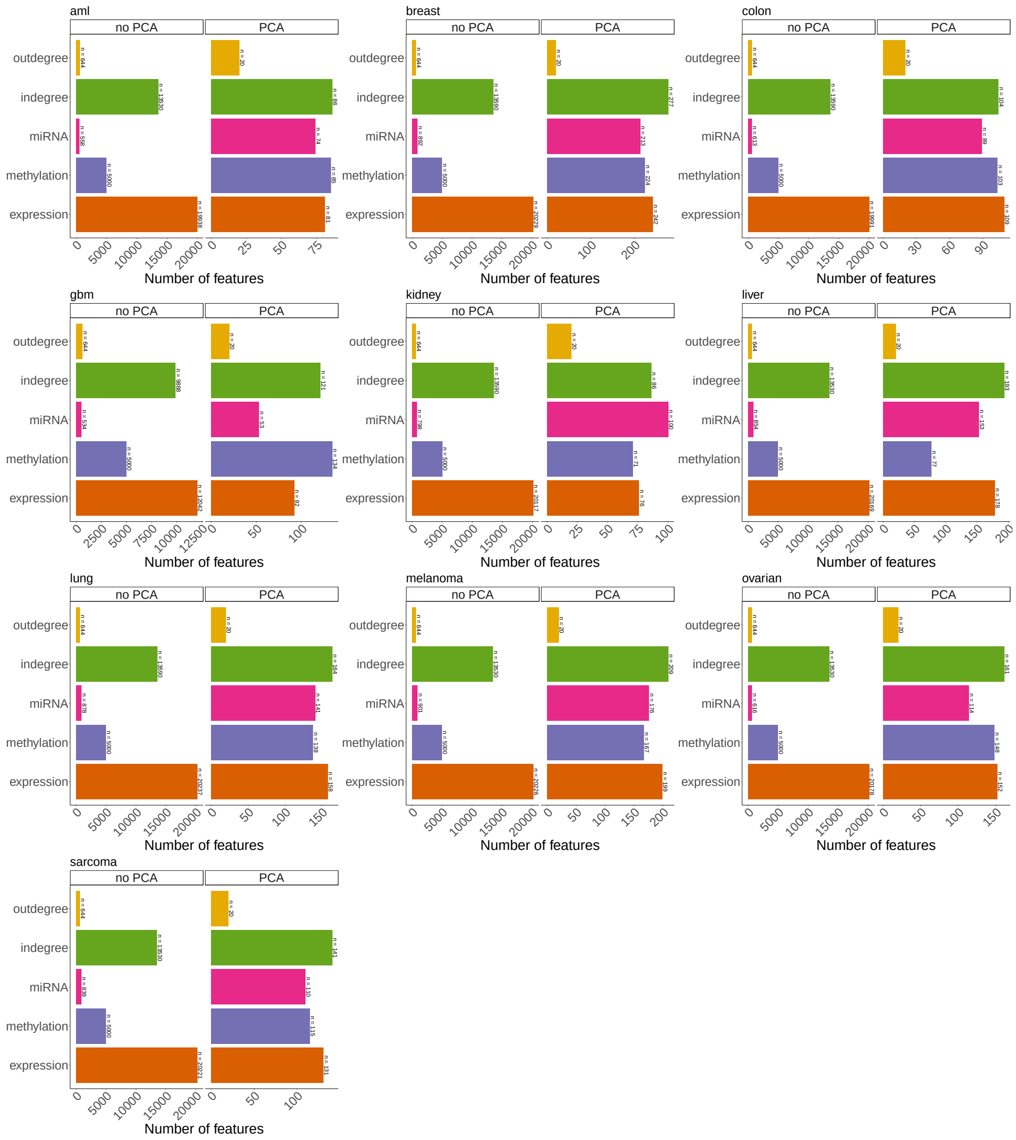


**Figure S3.** Comparison of data dimensions before and after performing PCA in ten cancer datasets from TCGA.

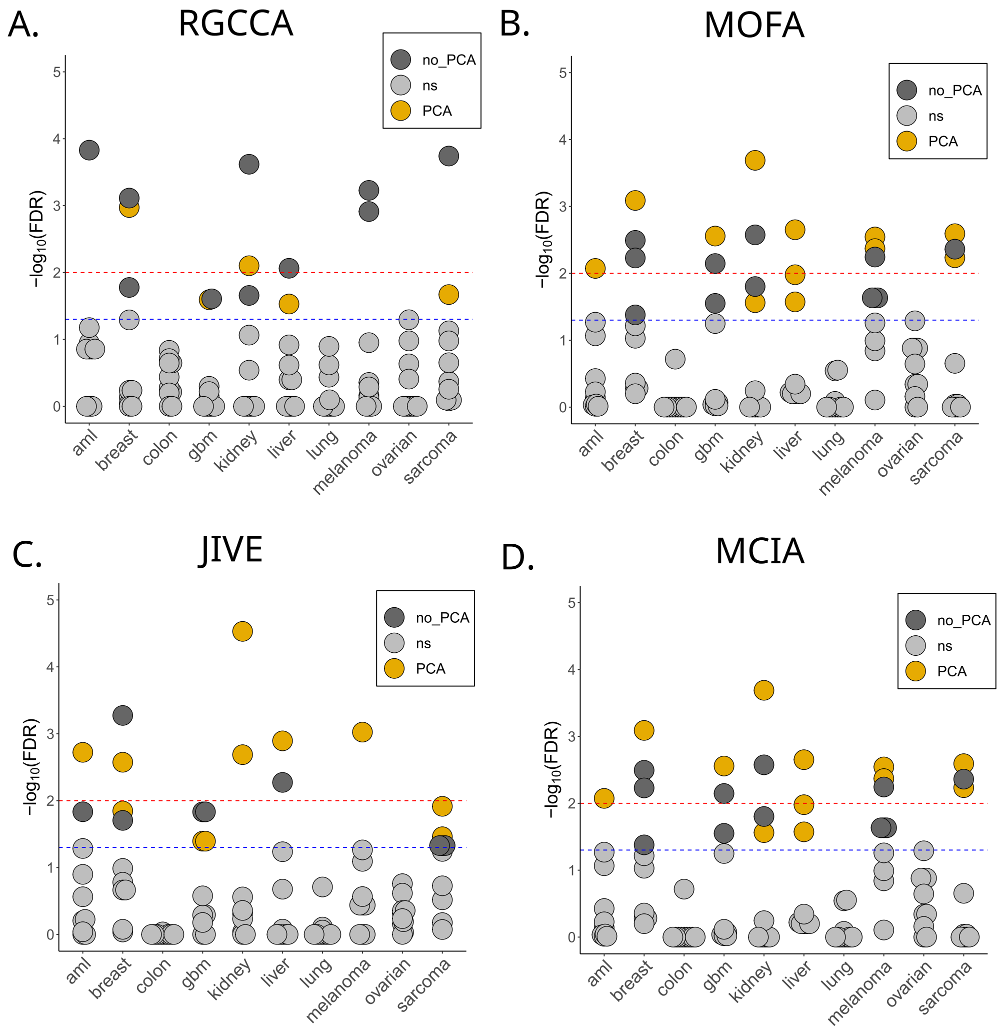


**Figure S4.** Comparison of factors association with survival with and without PCA in ten cancer datasets from TCGA using four different JDR tools: RGCCA (**A**), MOFA (**B**), JIVE (**C**) and MCIA (**D**).

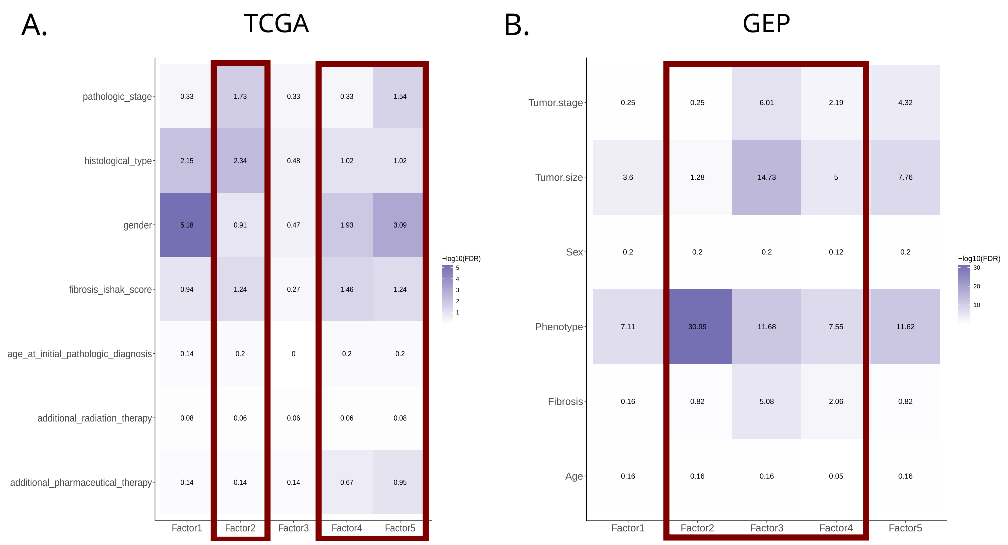


**Figure S5.** Association of MOFA factors with various clinical features in the TCGA liver (**A**) and GEP liver (**B**) datasets. Highlighted in red are the SAFs.

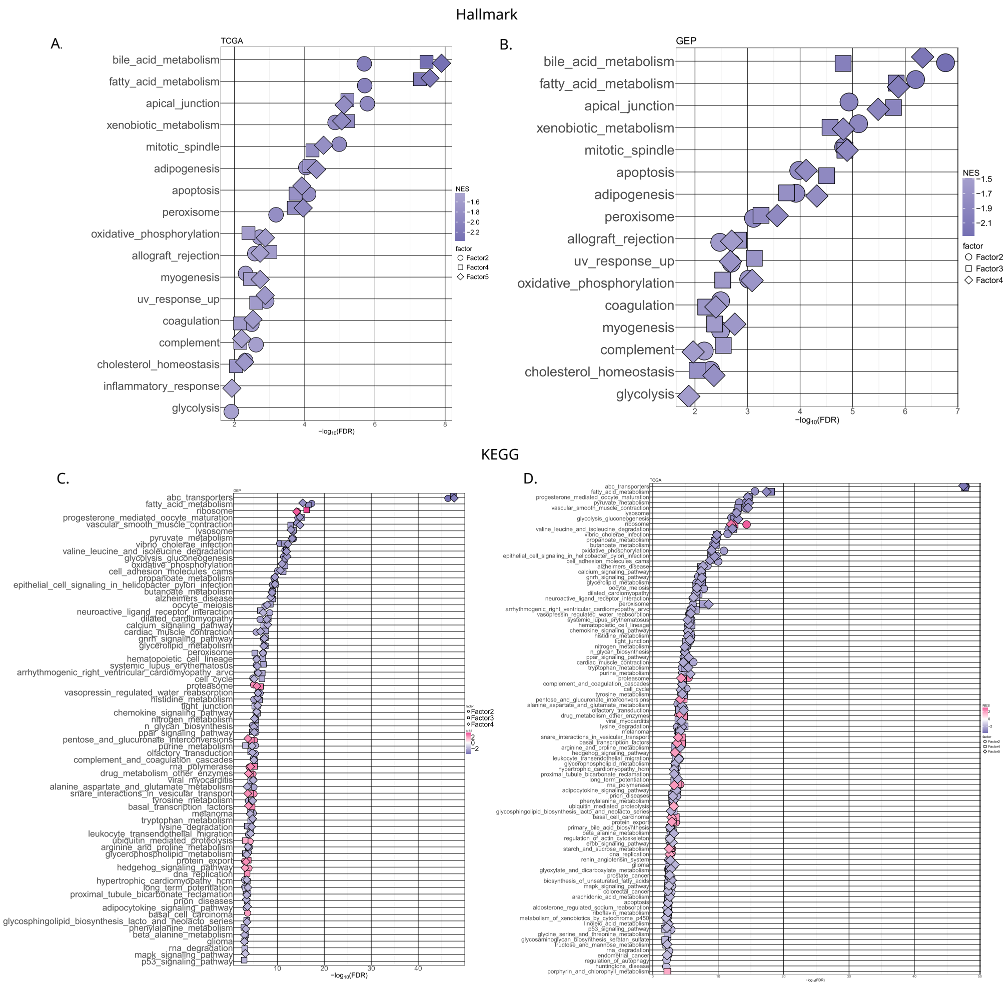


**Figure S6.** GSEA on indegree factor weights using the Hallmark (**A-B**) and KEGG (**C-D**) gene sets, showing all significant pathways with FDR ¡ 0.01

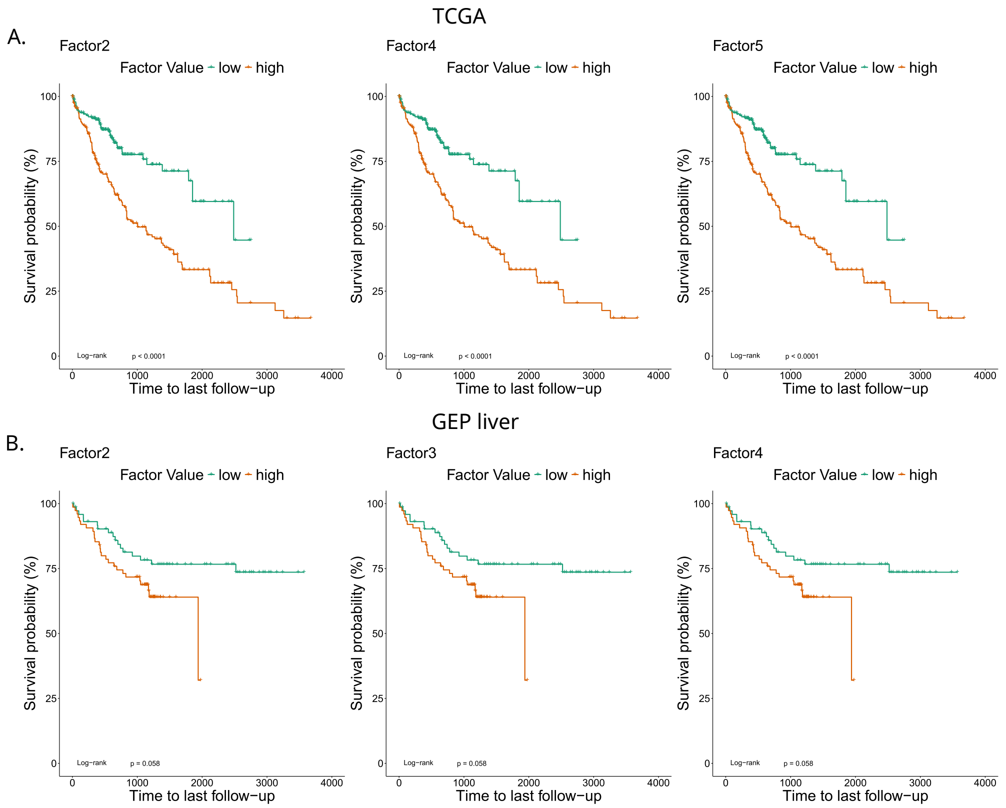


**Figure S7.** Kaplan-Meier curves when splitting the TCGA (**A**) and GEP (**B**) cohorts based on the median of MOFA survival associated factors.

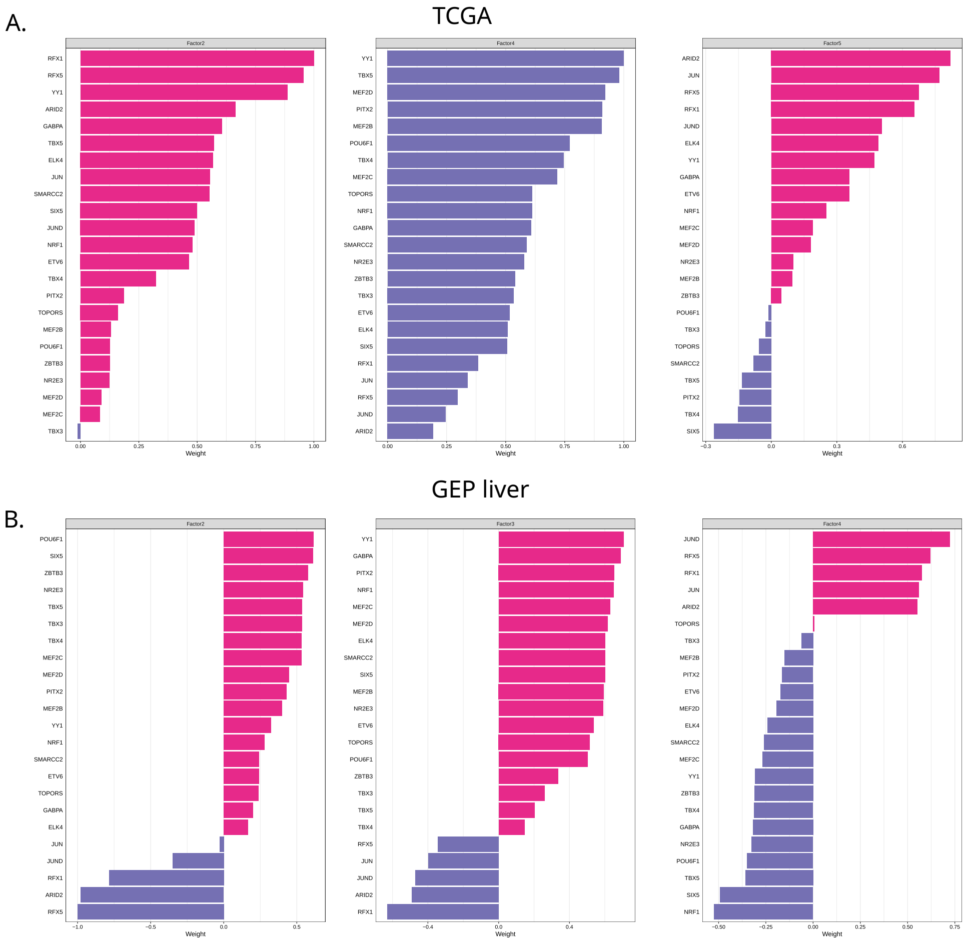


**Figure S8.** Weights of the 23 TFs shared between the TCGA (A) and GEP (B) datasets when selecting based on a absolute MOFA weight $\geq 0.5$

### Supplementary tables

**Table S1.** Number of samples used for each dataset for GRN inference, for JDR and for survival analysis.

**Table S2.** Summary of cox regression results of the association of MOFA factors with patient survival when using different seeds in ten TCGA cancer datasets.

**Table S3.** Summary of cox regression results for the association of MOFA factors with patient survival in a liver cancer dataset from GEPliver.
